# Functional equivalence of genome sequencing analysis pipelines enables harmonized variant calling across human genetics projects

**DOI:** 10.1101/269316

**Authors:** Allison A. Regier, Yossi Farjoun, David Larson, Olga Krasheninina, Hyun Min Kang, Daniel P. Howrigan, Bo-Juen Chen, Manisha Kher, Eric Banks, Darren C. Ames, Adam C. English, Heng Li, Jinchuan Xing, Yeting Zhang, Tara Matise, the NHLBI Trans-Omics for Precision Medicine (TOPMed) Program, Goncalo R. Abecasis, Will Salerno, Michael C. Zody, Benjamin M. Neale, Ira M. Hall

**Affiliations:** McDonnell Genome Institute, Washington University School of Medicine, St. Louis, MO, USA; Broad Institute of MIT and Harvard, Cambridge, MA, USA; Human Genome Sequencing Center, Baylor College of Medicine, Houston, TX, USA; Department of Biostatistics, University of Michigan, Ann Arbor, MI, USA; New York Genome Center, New York, NY, USA; DNAnexus Inc., Mountain View, CA, USA; Spiral Genetics, Seattle, WA, USA; Department of Genetics, Rutgers University, Piscataway, NJ, USA; Stanley Center for Psychiatric Research, Broad Institute of MIT and Harvard, Cambridge, MA, USA; Analytic and Translational Genetics Unit, Massachusetts General Hospital, Boston, MA, USA

## Abstract

Hundreds of thousands of human whole genome sequencing (WGS) datasets will be generated over the next few years to interrogate a broad range of traits, across diverse populations. These data are more valuable in aggregate: joint analysis of genomes from many sources increases sample size and statistical power for trait mapping, and will enable studies of genome biology, population genetics and genome function at unprecedented scale. A central challenge for joint analysis is that different WGS data processing and analysis pipelines cause substantial batch effects in combined datasets, necessitating computationally expensive reprocessing and harmonization prior to variant calling. This approach is no longer tenable given the scale of current studies and data volumes. Here, in a collaboration across multiple genome centers and NIH programs, we define WGS data processing standards that allow different groups to produce “functionally equivalent” (FE) results suitable for joint variant calling with minimal batch effects. Our approach promotes broad harmonization of upstream data processing steps, while allowing for diverse variant callers. Importantly, it allows each group to continue innovating on data processing pipelines, as long as results remain compatible. We present initial FE pipelines developed at five genome centers and show that they yield similar variant calling results – including single nucleotide (SNV), insertion/deletion (indel) and structural variation (SV) – and produce significantly less variability than sequencing replicates. Residual inter-pipeline variability is concentrated at low quality sites and repetitive genomic regions prone to stochastic effects. This work alleviates a key technical bottleneck for genome aggregation and helps lay the foundation for broad data sharing and community-wide “big-data” human genetics studies.

## Main text

Over the past few years, a wave of large-scale WGS-based human genetics studies have been launched by various institutes and funding programs worldwide, aimed at elucidating the genetic basis of a variety of human traits. These projects will generate hundreds of thousands of publicly available deep (>20x) WGS datasets from diverse human populations. Indeed, at the time of writing, >150,000 human genomes have already been sequenced by three NIH programs: NHGRI Centers for Common Disease Genomics^1^ (CCDG), NHLBI Trans-Omics for Precision Medicine^2^ (TOPMed), and NIMH Whole Genome Sequencing in Psychiatric Disorders^3^ (WGSPD). Systematic aggregation and co-analysis of these (and other) genomic datasets will enable increasingly well-powered studies of human traits, population history and genome evolution, and will provide population-scale reference databases that expand upon the groundbreaking efforts of the 1000 Genomes Project^4,5^, Haplotype Reference Consortium^6^, ExAC^7^ and GnomAD^8^.

Our ability as a field to harness these collective data to their full analytic potential depends on the availability of high quality variant calls from large populations of individuals. Accurate population-scale variant calling in turn requires joint analysis of all constituent raw data, where different batches have been aligned and processed systematically using compatible methods. Genome aggregation efforts are stymied by the distributed nature of human genetics research, where different groups routinely employ different alignment, data processing and variant calling methods. These methods often have comparable overall quality, but exhibit trivial incompatibilities that produce batch effects, limiting the utility of combined datasets. Prior exome/genome aggregation efforts have therefore been forced to obtain raw sequence data and re-perform upstream read alignment and data processing steps prior to joint variant calling^7,8^. These upstream steps are computationally expensive – representing as much as ~80% of the overall cost of WGS data analysis – and having to rerun them is inefficient. This computational burden will be increasingly difficult to bear as data volumes grow over coming years.

To help alleviate this burden and enable future genome aggregation efforts, we have forged a collaboration of major U.S. genome sequencing centers and NIH programs, and collaboratively defined data processing and file format standards to guide ongoing and future sequencing studies. Our approach focuses on the harmonization of upstream steps prior to variant calling, thus reducing trivial variability in core pipeline components while promoting the application of diverse and complementary variant calling methods – an area of much ongoing innovation. The guiding principle is the concept of “functional equivalence” (FE). We define FE to be a shared property of two pipelines that can be run independently on the same raw WGS data to produce two output files that, upon analysis by the same variant caller(s), produce virtually indistinguishable genome variation maps. A key question, of course, is where to draw the FE threshold. There is no one answer; at minimum, we advise that data processing pipelines should introduce much less variability in a single DNA sample than independent WGS replicates of DNA from the same individual.

Towards this goal, we defined a set of required and optional data processing steps and file format standards (Fig. 1; see GitHub page^9^ for details). We focus here on WGS data analysis, but these guidelines are equally suitable for exome sequencing. These standards are founded in extensive prior work in the area of read alignment^10^, sequence data analysis^5,11-17^ and compression^11,18^, and more broadly in WGS analysis best practices employed at our collective institutes, and worldwide. Notable features of the data processing standard include alignment with BWA-MEM^10^, adoption of a standard GRCh38 reference genome with alternate loci^4,19,20^, and improved duplicate marking. File format standards include a 4-bin base quality scheme, CRAM compression^18^ and restricted tag usage, which in combination reduced file size >3-fold (from 54 to 17 Gb for a 30X WGS and from 38 to 12 Gb for a 20X WGS). This in turn reduces data storage costs and increases transfer speeds, facilitating data access and sharing.

**Figure 1.**
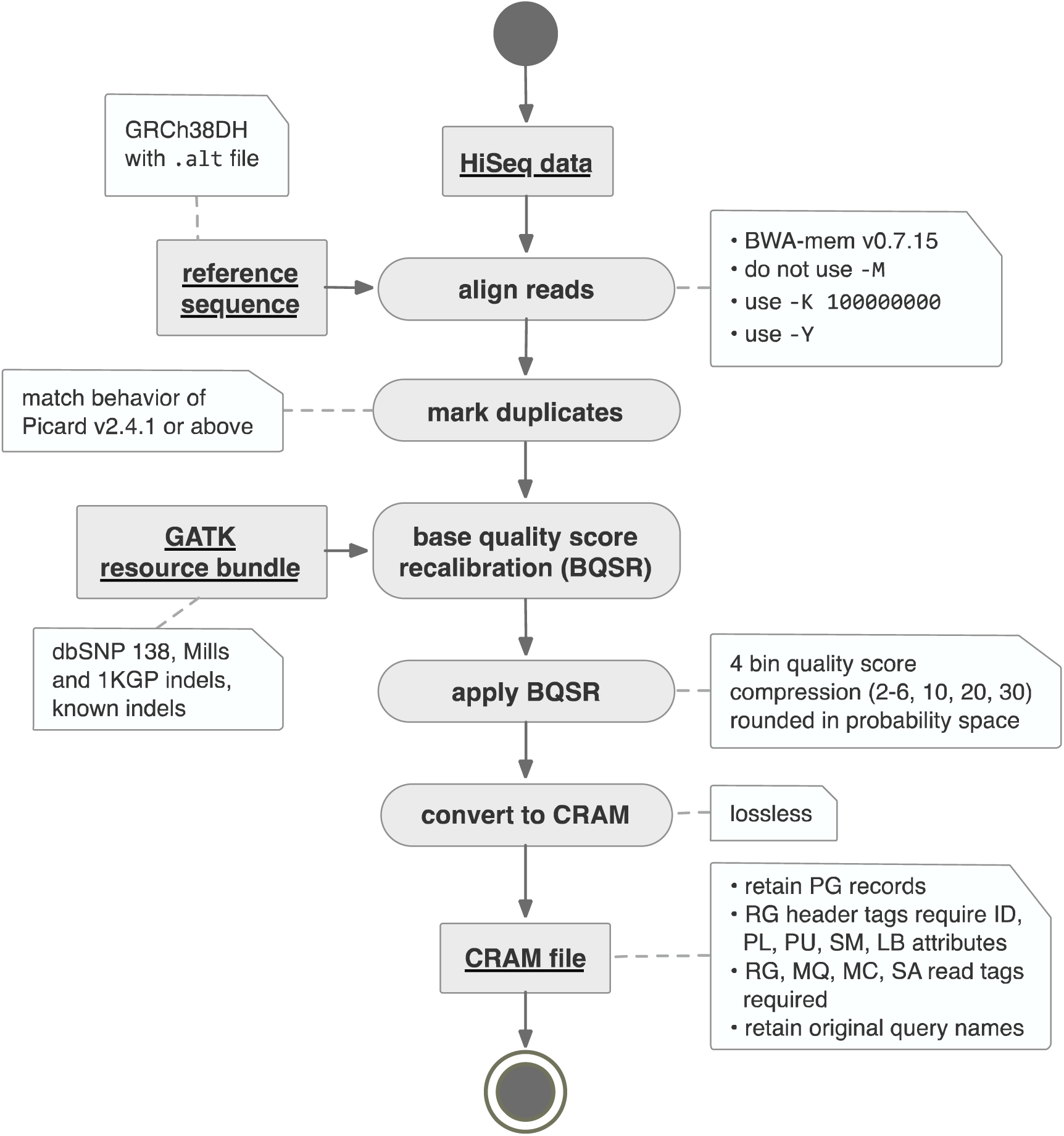
We defined a series of required and allowed processing steps that provide flexibility in pipeline implementation while keeping variation between pipelines at a minimum. Reads must be aligned to a specific reference genome using a minimum version of the BWA-MEM aligner. Algorithms for marking duplicates and recalibrating base quality scores are more flexible and vary somewhat between centers. Compression of quality scores into four bins saves storage and file transfer costs, while maintaining acceptable accuracy and sensitivity.

We implemented initial versions of these pipelines at each of the five participating centers, including the four CCDGs as well as the TOPMed Informatics Resource Core, and serially tested and modified them based on alignment statistics (**Supplementary Table 1)** and variant calling results from a 14-genome test set, using GATK^21^ for single nucleotide variants (SNVs) and small insertion/deletion (indel) variants, and LUMPY^22^ for structural variants (SVs), with data contributed from each center (see **Methods**). These 14 datasets have diverse ancestry and are composed of well-studied samples from the 1000 Genomes Project^4^, including 4 independently-sequenced replicates of NA12878 (CEPH) and 2 replicates of NA19238 (Yoruban). We tested pairwise variability in SNV, indel and SV callsets generated separately from each of the five pipelines, before and after harmonization, as compared to variability between WGS data replicates (Fig. 2). As expected, pipelines used by centers prior to harmonization effort exhibit strong levels of variability, especially among SV callsets. Most importantly, variability between harmonized pipelines (mean 0.4%, 1.8%, and 1.1% discordant for SNVs, indels, and SVs, respectively) is an order of magnitude lower than between replicate WGS datasets (mean 7.1%, 24.0%, and 39.9% discordant). Note that absolute levels of discordance are somewhat high in this analysis because we performed per-sample variant calling and included all genomic regions, with minimal variant filtering. All pipelines show similar levels of sensitivity and accuracy based on Genome in a Bottle (GiaB) calls for NA12878^23^ (Supplementary Fig. 1).

**Figure 2.**
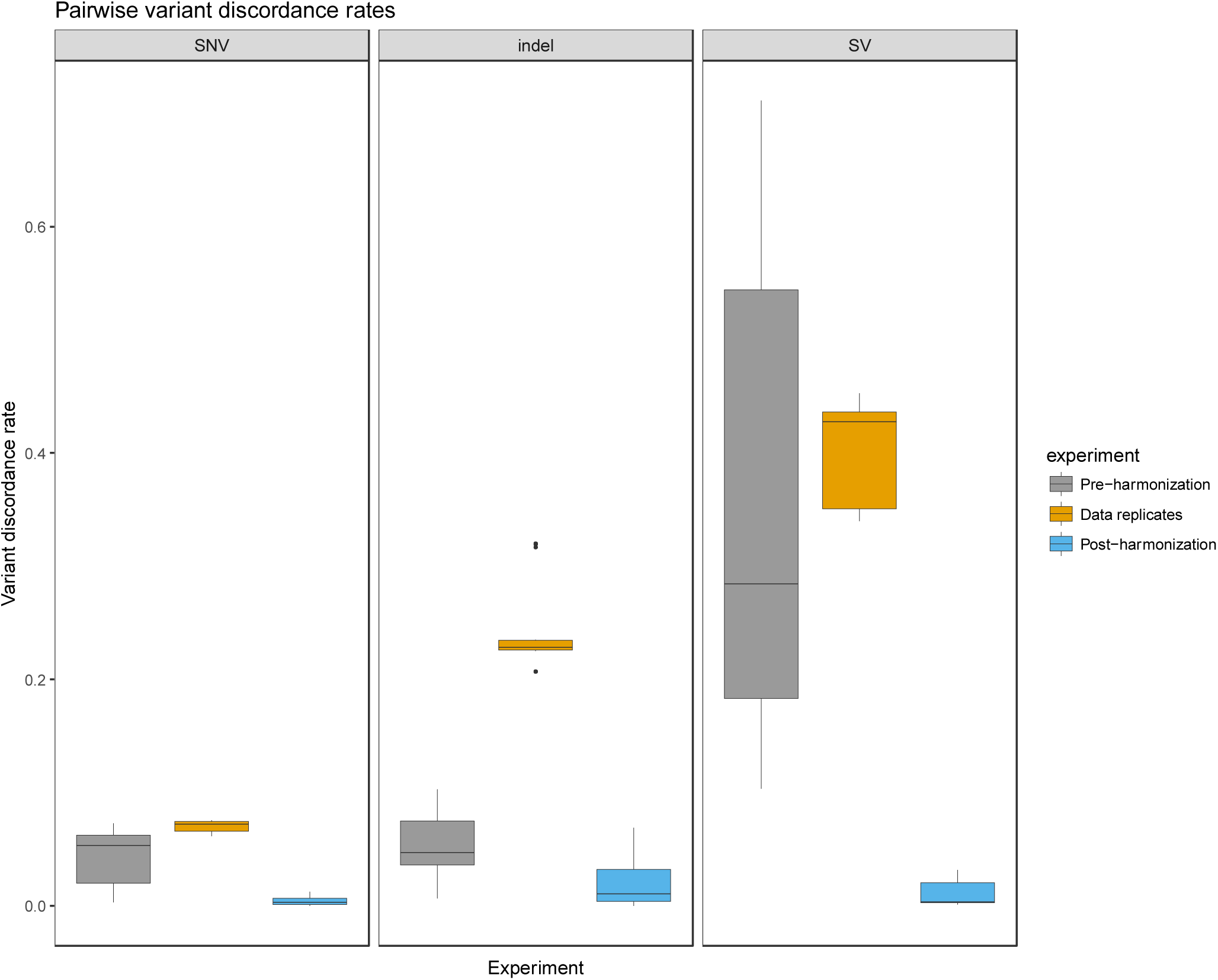
Pairwise variant discordance rates were calculated between pipelines from each of five centers (pre-harmonization and post-harmonization) as well as between independent sequencing replicates of the same individuals processed by the same pipeline (data replicates). From left, single nucleotide (SNV) and small insertion/deletion (indel) variants were detected with GATK, and structural variants (SV) with LUMPY. The pre- and post-harmonization comparisons include 14 independently sequenced samples. The data replicate comparisons include four replicates of NA12878 and two replicates of NA19238. Note that the extremely high levels of discordance for SVs pre-harmonization are largely due to variable use of “decoy” sequences in the reference genomes used by the different centers.

We next applied the final pipeline versions to an independent set of 100 genomes comprising 8 trios from the 1000 Genomes Project^4,5^ and 19 quads from the Simons Simplex Collection^24^, and generated separate 100-genome GATK and LUMPY callsets using data from each of the five pipelines. Considering all five callsets in aggregate, the vast majority of GATK variants (97.2%) are identified in data from all five pipelines, with only 1.74% unique to a single pipeline and 1.02% in various minor subsets. Mean pairwise SNV concordance rates are in the range of 99.0-99.9% over all sites and comparisons, and Mendelian error rates are ~0.3% at concordant sites, and ~22-24% at discordant sites (Fig. 3). Indel and SV concordance rates are lower – as expected given that these variants are more difficult to map and genotype precisely. Pairwise SNV concordance rates are substantially higher in GiaB high confidence genomic regions comprised predominantly of unique sequence (SNV concordance: 99.7-99.9%; 72% of genome) than in difficult-to-assess regions laden with segmental duplications and high copy repeats (SNV concordance: 92-99%; 8.5% of genome; **see Methods**). Indeed, 58% of discordant SNV calls are found in the 8.5% most difficult to analyze subset of the genome. Furthermore, the mean quality score of discordant SNV sites are only 0.5% as high as the mean score of concordant SNV sites (16.4% for indels and 90.0% for SVs) (Supplementary Fig. 2). This suggests that many discordant sites are either false positive calls or represent sites that are difficult to measure robustly with current methods. Differences between pipelines are roughly symmetric, with all pipelines achieving similarly low levels of performance at discordant sites, as based on pairwise discordance rates and Mendelian error rates (Supplementary Fig. 3), further suggesting that most discordant calls are due to stochastic effects at sites with borderline levels of evidence. We note that there are some center-specific sources of variability due to residual differences in BQSR models and alignment filtering methods, but that these affect only a trivial fraction of variant calls.

**Figure 3.**
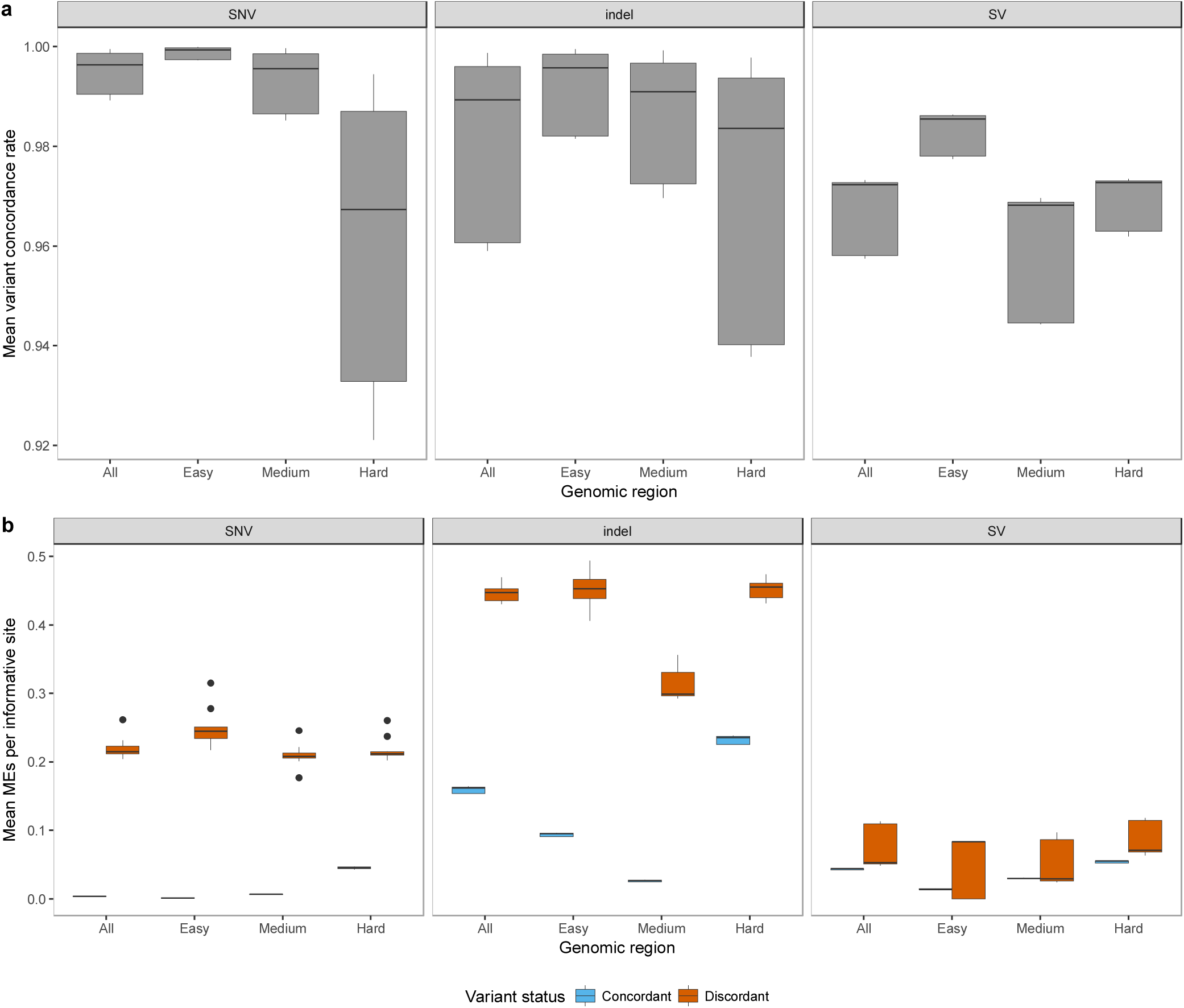
Variant concordance and Mendelian error (ME) rates for different variant classes and genomic regions using 100 samples, including 8 trios from the 1000 Genomes Project and 19 quads from the Simons Simplex Collection. **(a)** Variant concordance rates were calculated from pairwise comparisons across five pipelines for 100 samples. **(b)** Mendelian error rates were calculated using informative sites in 44 parent-offspring trios, for variants classified as concordant and discordant in pairwise comparisons between five pipelines.

Here, we have described a simple yet effective approach for harmonizing data processing pipelines through the concept of “functional equivalence”. This work resolves a key source of batch effects in sequencing data from different genome centers, and thus alleviates a bottleneck for data sharing and collaborative analysis within and among large-scale human genetics studies. Our approach also facilitates accurate comparison to variant databases; researchers that want to analyze their sample(s) against major datasets such as gnomAD, TOPMed, or CCDG should adopt these standards in order to avoid artifacts caused by non-FE sample processing. Of course, other challenges remain, such as batch effects from library preparation and sequencing, and persistent regulatory hurdles. Nevertheless, we envision that it will be possible to robustly generate increasingly large genome variation maps and shared annotation resources from these and other programs over the next few years, from diverse groups and analysis methods. Ultimately, we hope that international efforts such as Global Alliance for Genomics & Health (GA4GH) will adopt and extend these guidelines to help integrate research and medical genomes worldwide.

## Acknowledgments

We thank NHGRI and NHLBI program staff for supporting this effort. This work was funded by NHGRI CCDG awards to Washington University in St. Louis (UM1 HG008853), Broad Institute of MIT and Harvard (UM1 HG008895), Baylor College of Medicine (UM1 HG008898), and the New York Genome Center (UM1 HG008901), the NHGRI GSP coordinating center (U24 HG008956), and an NHLBI TOPMed Informatics Research Center award to the University of Michigan (3R01HL-117626-02S1) as well as grants to B.M.N. (U01 HG00908, R01 MH107649), H.M.K (1 R21 HL133758-01, 1 U01 HL137182-01) and G.R.A. (4 R01 HL117626-04)

## Methods

### Dataset selection

For initial testing, we selected 14 whole genome sequencing datasets based on the following criteria: (1) they include samples of diverse ancestry, including CEPH (NA12878, NA12891, NA12892), Yoruban (NA19238), Luhya (NA19431), and Mexican (NA19648); (2) they were sequenced at multiple different genome centers to deep coverage (>20X) using Illumina HiSeq X technology; (3) they include replicates of multiple samples, including 2 of NA19238 (Yoruban) and 4 of NA12878 (CEPH); (4) they include the extremely well-studied NA12878 genome, for which much ancillary data exists, and (5) they were open access, readily accessible and shareable among the consortium sites. For subsequent characterization of the finalized pipelines, we selected an independent set of 100 samples composed of 8 open-access trios of diverse ancestry from the 1000 Genomes project – including CEPH (NA12878, NA12891, NA12892), Yoruban (NA19238, NA19239, NA19240), Southern Han Chinese (HG00512, HG00513, HG00514), Puerto Rican (HG00731, HG00732, HG00733), Colombian (HG01350, HG01351, HG01352), Vietnamese (HG02059, HG02060, HG02061), Gambian (HG02816, HG02817, HG02818), and Caucasian (NA24143, NA24149, NA24385) – and 19 quads from the Simons Simplex Collection^24^.

### Downsampling data replicates

To eliminate coverage differences as a contributor to variation between sequencing replicates of the same sample (4 replicates of NA12878 and 2 replicates of NA19238), the data replicates were downsampled to match the lowest coverage sample. To obtain initial coverage, all replicates were aligned to a build 37 reference using speedseq^14^ (v 0.1.0). Mean coverage for each BAM file was calculated using the Picard CollectWgsMetrics tool (v2.4.1)^12^. For each sample, a downsampling ratio was calculated using the lowest coverage as the numerator and the sample’s coverage as the denominator. This ratio was used as the PROBABILITY parameter for the Picard DownsampleSam tool, along with RANDOM_SEED=1 and STRATEGY=ConstantMemory. The resulting BAM was converted to FASTQ using the script bamtofastq.py from the speedseq repository.

### Alignment and data processing pipelines - WashU, pre- and post-harmonization

The pre-harmonization pipeline aligns reads to the GRCh37-lite reference using speedseq (v0.1.0)^14^. This includes alignment using bwa (v0.7.10-r789)^10^, duplicate marking using samblaster (v0.1.22)^13^, and sorting using sambamba (v0.5.4)^16^.

The post-harmonization pipeline aligns each read group separately to the GRCh38 reference using bwa-mem (v0.7.15-r1140) with the parameters ‘-K 100000000 -p -Y’. MC and MQ tags are added using samblaster (v0.1.24) with the parameters ‘-a --addMateTags’. Read group BAM files are merged together with ‘samtools merge’ (v1.3.1-2). The resulting file is name-sorted with ‘sambamba sort -n’ (v0.6.4). Duplicates are marked using Picard MarkDuplicates (v2.4.1) with the parameter ‘ASSUME_SORT_ORDER=queryname’, then the results are coordinate sorted using ‘sambamba sort’. A base quality recalibration table is generated using GATK BaseRecalibrator (v3.6) with knownSites files (dbSNP138, Mills and 1kg indels, and known indels) from the GATK resource bundle (https://console.cloud.google.com/storage/browser/genomics-public-data/resources/broad/hg38/v0) and parameters ‘--preserve_qscores_less_than 6 -dfrac .1 -nct 4 -L chr1 -L chr2 -L chr3 -L chr4 -L chr5 -L chr6 -L chr7 -L chr8 -L chr9 -L chr10 -L chr11 -L chr12 -L chr13 -L chr14 -L chr15 -L chr16 -L chr17 -L chr18 -L chr19 -L chr20 -L chr21 -L chr22’. The base recalibration table is applied using GATK PrintReads with the parameters ‘-preserveQ 6 -BQSR “${bqsrt}” -SQQ 10 -SQQ 20 -SQQ 30 -- disable_indel_quals’. Finally, the output is converted to CRAM using ‘samtools view’.

### Alignment and data processing pipelines – Broad, pre- and post-harmonization Pre harmonization

-Align with bwa-mem v0.7.7-r441: bwa mem –M –t 10 –p GRCh37.fasta

-Merge aligned bam with the original unaligned bam and sort with Picard 2.8.3: MergeBamAlignment ADD_MATE_CIGAR=true ALIGNER_PROPER_PAIR=false UNMAP_CONTAMINANT_READS=false SORT_ORDER=coordinate

-Mark duplicates with Picard 2.8.3: MarkDuplicates

-Find target indels to fix with GATK 3.4-g3c929b0: CreateRealignerTargets –known dbSnp.138.vcf –known mills.vcf –known 1000genome.vcf

-Fix indel alignments with GATK 3.4-g3c929b0: –known dbSnp.138.vcf –known mills.vcf –known 1000genome.vcf

-Create recalibration table using GATK 3.4-g3c929b0: RecalibrateBaseQuality –knownSites dbSnp.138.vcf using –known dbSnp.138.vcf –known mills.vcf –known 1000genome.vcf

-Apply base recalibration using GATK 3.4-g3c929b0: PrintReads –disable_indel_quals – emit_original_quals

### Post harmonization

- Align with bwa-mem 0.7.15.r1140: bwa mem -K 100000000 -p -v 3 -t 16 –Y GRCh38.fasta

-Merge aligned bam with the original unaligned bam with Picard 2.16.0: MergeBamAlignment EXPECTED_ORIENTATIONS=FR ATTRIBUTES_TO_RETAIN=X0 ATTRIBUTES_TO_REMOVE=NM ATTRIBUTES_TO_REMOVE=MD REFERENCE_SEQUENCE=${ref_fasta} PAIRED_RUN=true SORT_ORDER="unsorted CLIP_ADAPTERS=false MAX_INSERTIONS_OR_DELETIONS=-1 PRIMARY_ALIGNMENT_STRATEGY=MostDistant UNMAPPED_READ_STRATEGY=COPY_TO_TAG ALIGNER_PROPER_PAIR_FLAGS=true UNMAP_CONTAMINANT_READS=true

-ADD_PG_TAG_TO_READS=false

-Mark duplicates with Picard 2.16.0: MarkDuplicates ASSUME_SORT_ORDER="queryname"

-Sort with Picard 2.16.0: SortSam SortOrder=coordinate

-Create BQSR table using GATK 4.beta.5: BaseRecalibrator

-knownSites dbSnp.138.vcf using –known dbSnp.138.vcf –known mills.vcf –known 1000genome.vcf

-Apply recalibration using GATK 4.beta.5:

ApplyBQSR -SQQ 10 -SQQ 20 -SQQ 30

-Convert output to cram with SamTools v 1.3.1: samtools view -C -T GRCh38.fasta

### Alignment and data processing pipelines – Baylor, pre & post harmonization

In the HGSC pre-harmonized WGS protocol (https://github.com/HGSC-NGSI/HgV_Protocol_Descriptions/blob/master/hgv_resequencing.md), reads are mapped to the GRCh37d reference with bwa-mem (v0.7.12), samtools (v1.3) fixmate, sorting and duplicate marking with sambamba (v0.5.9), base recalibration and realignment with GATK (v3.4.0), and the quality scores are binned and tags removed with bamUtil squeeze (v1.0.13). Multiplexed samples follow the same steps up through sorting and duplicate marking, resulting in sequencing-event BAMs. The BAMs are merged and duplicates marked using sambamba (v0.5.9), followed by the recalibration, realignment and binning described above.

The HGSC harmonized WGS protocol (https://github.com/HGSC-NGSI/HgV_Protocol_Descriptions/blob/master/hgv_ccdg_resequencing.md) aligns each read group to the GRCh38 reference using bwa-mem (0.7.15) with the parameters ‘-K 100000000 -Y’. MC and MQ tags are added using samblaster (v0.1.24) with the parameters ‘-a --addMateTags’. The resulting file is name-sorted with ‘sambamba sort -n’ (v0.6.4). Duplicates are marked using Picard MarkDuplicates (v2.4.1) with the parameter ‘ASSUME_SORT_ORDER=queryname’, then the results are coordinate-sorted using ‘sambamba sort’. For multiplexed samples, these sequence-event BAMs are then merged with sambamba (v0.6.4) merge, name sorted, duplicate marked and coordinate-sorted with the same tools as above. A base quality recalibration table is generated using GATK BaseRecalibrator (v3.6) with knownSites files (dbSNP138, Mills and 1kg indels, and known indels) from the GATK resource bundle (https://console.cloud.google.com/storage/browser/genomics-public-data/resources/broad/hg38/v0) and parameters ‘--preserve_qscores_less_than 6 -dfrac .1 -nct 4 -L chr1 -L chr2 -L chr3 -L chr4 -L chr5 -L chr6 -L chr7 -L chr8 -L chr9 -L chr10 -L chr11 -L chr12 -L chr13 -L chr14 -L chr15 -L chr16 -L chr17 -L chr18 -L chr19 -L chr20 -L chr21 -L chr22’. The base recalibration table is applied using GATK PrintReads with the parameters ‘-preserveQ 6 -BQSR “${bqsrt}” -SQQ 10 -SQQ 20 -SQQ 30 -- disable_indel_quals’. Finally, the output is converted to CRAM using ‘samtools view’.

### Alignment and data processing pipelines – NYGC, pre- and post-harmonization

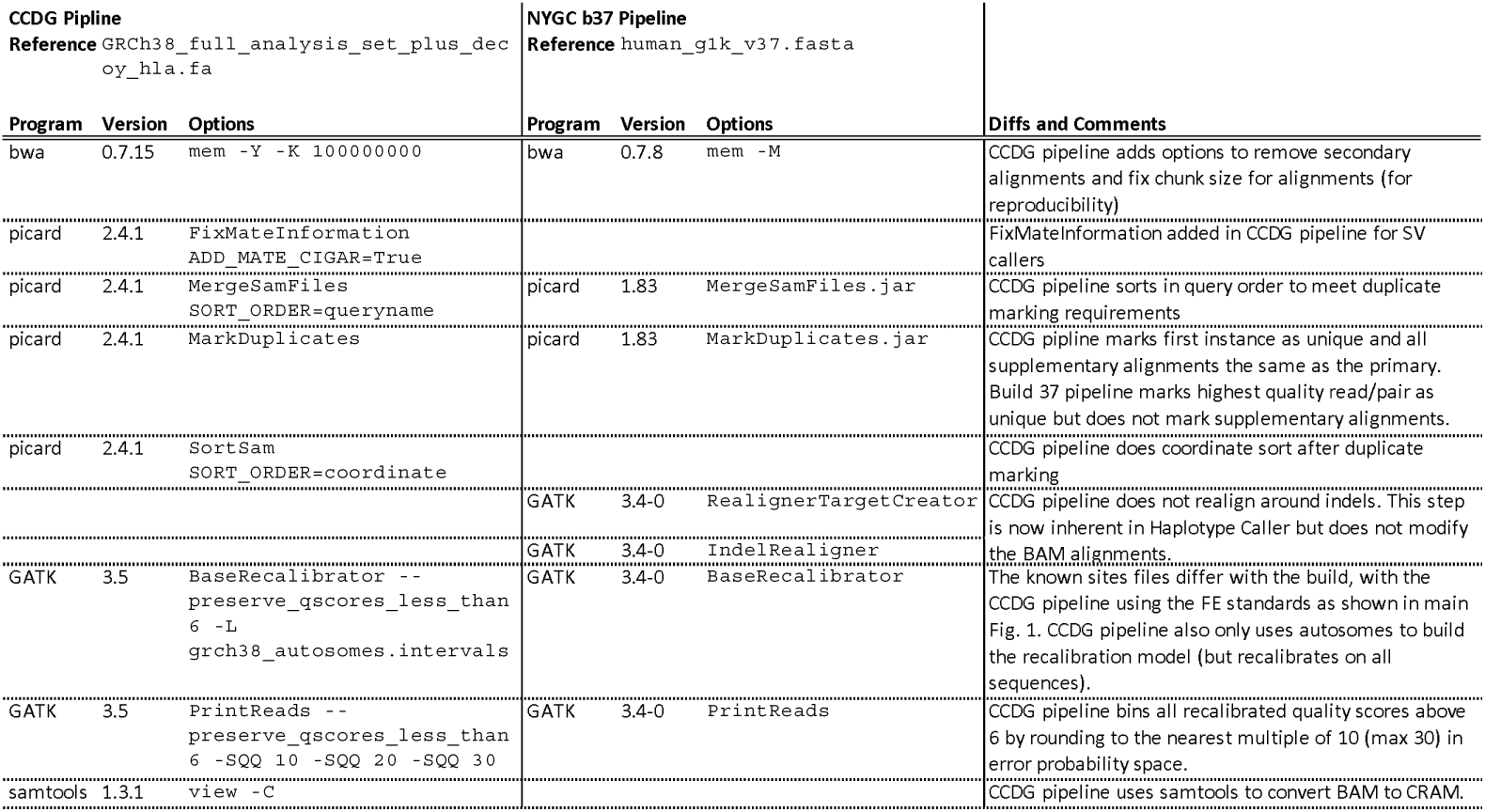

### Alignment and data processing pipelines – Michigan, pre- and post-harmonization

The pre-harmonization pipeline aligns reads using default options in the GotCloud alignment pipeline^15^ available at https://github.com/statgen/gotcloud. It aligns the sequence reads to GRCh37 reference with decoy sequences used in 1000 Genomes. The raw sequence was aligned using bwa mem (v0.7.13- r1126)^10^, and sorted by samtools (v1.3.1). The duplicate marking and base quality recalibration were performed jointly using bamUtil dedup [ref – same as GotCloud] (v1.0.14).

The post-harmonization pipeline procedure (described in https://github.com/statgen/docker-alignment) first aligns each read group to the GRCh38 reference using bwa-mem (v0.7.15-r1140) with the parameters ‘-K 100000000 -Y -R [read_group_id]’. To add MC and MQ tags, samblaster (v0.1.24) was used with the parameters ‘-a --addMateTags’. Each BAM file corresponding to a read group is sorted by genomic coordinate using ‘samtools sort’ (v1.3.1), and merged together using ‘samtools merge’ (v1.3.1). Duplicate marking and base quality recalibration were performed jointly using bamUtil dedup_lowmem (v1.0.14). with and parameters ‘--allReadNames –binCustom –binQualS 0:2,3:3,4:4,5:5,6:6,7:10,13:20,23:30,33:40 --recab --refFile [reference_fasta_file] --dbsnp [dbsnp_b142_vcf_file] --in [input_bam] –out -.ubam’ and the piped output (in uncompressed BAM format) is convered into s a CRAM file using samtools view.

### Calculation of Alignment Statistics

A total of 184 alignment statistics were generated for all standardized CRAM files from each center with AlignStats software. Results include metrics for both the entire CRAM file and for the subset of read-pairs with at least one read mapping to the autosome or sex chromosomes. We examined all metrics across the five CRAMs for each of the 15 samples to ensure that any differences were consistent with the various options allowed in the functional equivalence specification. Supplementary Table 1 provides examples of these metrics, and full description of all metrics can be found online (https://github.com/jfarek/alignstats.

### Variant calling for the 14-sample analysis

SNPs and indels were called for each center’s CRAM/BAM files using GATK^21^ version 3.5-0-g36282e4 HaplotypeCaller with the following parameters:

-rf BadCigar

--genotyping_mode DISCOVERY

--standard_min_confidence_threshold_for_calling 30

--standard_min_confidence_threshold_for_emitting 0

For the pre-standardization files, the 1000 genomes phase 3 reference sequence from the GATK reference bundle ftp://ftp.broadinstitute.org/pub/svtoolkit/reference_metadata_bundles/1000G_phase3_25Jan2015.tar.gz was used. For the post-standardization files, the 1000 Genomes Project version of GRCh38DH (http://ftp.1000genomes.ebi.ac.uk/vol1/ftp/technical/reference/GRCh38_reference_genome/) was used.

Structural variants (SVs) were called for each center’s CRAM/BAM files using lumpy^22^ and svtools (https://github.com/hall-lab/svtools). First, split reads and reads with discordant insert sizes or orientations were extracted from the CRAM/BAM files using extract-sv-reads in the docker image halllab/extract-sv-reads@sha256:192090f72afaeaaafa104d50890b2fc23935c8dc98988a9b5c80ddf4ec50f70c using the following parameters:

--input-threads 4 -e –r

Next, SV calls were made using lumpyexpress (https://github.com/arq5x/lumpy-sv) from the docker image halllab/lumpy@sha256:59ce7551307a54087e57d5cec89b17511d910d1fe9fa3651c12357f0594dcb07 with the -P parameter as well as -x to exclude regions contained in the BED file exclude.cnvnator_100bp.GRCh38.20170403.bed (exclude.cnvnator_100bp.112015.bed for pre-standardization samples). Both exclude files are available in https://github.com/hall-lab/speedseq/tree/master/annotations

Finally, the SV calls were genotyped using svtyper [https://github.com/hall-lab/svtools/tree/master/svtools/bin/svtyper] from the docker image halllab/svtyper@sha256:21d757e77dfc52fddeab94acd66b09a561771a7803f9581b8cca3467ab7ff94a

### Defining “easy”, “medium” and “hard” genomic regions

The reference genome sequence is not uniformly amenable to analysis – some regions with high amounts of repetitive sequence are difficult to align and prone to misleading analyses, while other regions comprised of mostly unique sequence can be more confidently interpreted. To gain a better understanding of how pipeline concordance differs by region, we divided the reference sequence into three broad categories. The “easy” genomic regions consist of the GiaB gold standard high confidence regions, lifted over to build 38. The “hard” regions consist of centromeres (https://www.ncbi.nlm.nih.gov/projects/genome/assembly/grc/human/data/38/Modeled_regions_for_GRCh38.tsv), microsatellite repeats (satellite entries from http://hgdownload.soe.ucsc.edu/goldenPath/hg38/bigZips/hg38.fa.out.gz), low complexity regions (https://github.com/lh3/varcmp/raw/master/scripts/LCR-hs38.bed.gz), and windows determined to have high copy number (more than 12 copies per genome across 409 samples). Any regions overlapping GiaB high confidence regions are removed from the set of hard regions. All remaining regions are classified as “medium”.

### Cross-center variant comparisons for the 14-sample analysis

The VCF files produced by GATK for both the pre- and post-standardization experiments were compared using hap.py[https://github.com/Illumina/hap.py] from the docker image pkrusche/hap.py:v0.3.9 using the --preprocess-truth parameter.

The four data replicates of NA12878 were compared to the NA12878 gold standards (ftp://ftp-trace.ncbi.nlm.nih.gov/giab/ftp/release/NA12878_HG001/NISTv2.19/NISTIntegratedCalls_14datasets_131103_allcall_UGHapMerge_HetHomVarPASS_VQSRv2.19_2mindatasets_5minYesNoRatio_all_nouncert_excludesimplerep_excludesegdups_excludedecoy_excludeRepSeqSTRs_noCNVs.vcf.gz in the regions defined by ftp://ftp-trace.ncbi.nlm.nih.gov/giab/ftp/release/NA12878_HG001/NISTv2.19/union13callableMQonlymerged_addcert_nouncert_excludesimplerep_excludesegdups_excludedecoy_excludeRepSeqSTRs_noCNVs_v2.19_2mindatasets_5minYesNoRatio.bed.gz) to obtain sensitivity and precision measurements. The post-standardization VCFs were first lifted over to GRCh37 using the Picard LiftoverVcf tool (v2.9.0) and the chain files hg38ToHg19.over.chain.gz and hg19ToGRCh37.over.chain.gz downloaded from here: http://crossmap.sourceforge.net/#chain-file. To reduce artifacts from the liftover that negatively impacted sensitivity, the gold standard files were lifted over to the build 38 reference and back to build 37, excluding any variants that didn’t lift over in both directions.

Values for sensitivity (METRIC.Recall) and precision (METRIC.Precision) were parsed out of the *.summary.csv file produced by hap.py for each comparison, using only variants with the PASS filter value set.

The downsampled data replicates of NA12878 and NA19238 aligned by the same center were compared to each other in a pairwise fashion. Pairwise comparisons between centers were performed for each non-downsampled aligned file. The variant discordance rates between pairs were calculated using the true positive, true negative, and false positive counts from the *.extended.csv output file from hap.py (TRUTH.FN + QUERY.FP)/(TRUTH.TP + TRUTH.FN + QUERY.FP). The rates reported are only for PASS variants but across the whole genome.

The VCF files of SVs produced by lumpy and svtyper were converted to BEDPE using the command ‘svtools vcftobedpe’ from the docker container halllab/svtools@sha256:f2f3f9c788beb613bc26c858f897694cd6eaab450880c370bf0ef81d85bf8d45 The coordinates are padded with 1 bp on each side to be compatible with bedtools pairtopair. The pairwise comparisons are performed using the bedtools pairtopair command (version 2.23.0), then summarized using a python script (compare_single_sample_based_on_strand.py in https://github.com/CCDG/Pipeline-Standardization). The variant discordance rates between pairs are calculated with the following formula: (discordant + 0-only + 1-only + discordant_discordant_type)/(match + discordant + match_discordant_type + discordant_discordant_type + 0-only + 1-only).

### Variant calling for 100-sample analysis

SNPs and indels were called using the GATK best practices pipeline, including per-sample variant discovery using HaplotypeCaller with the following parameters: ‘-ERC GVCF -GQB 5 -GQB 20 -GQB 60 -variant_index_type LINEAR -variant_index_parameter 128000’. Next, GVCFs from all 100 samples were merged with GATK CombineGVCFs. Genotypes were refined with GATK GenotypeGVCFs with the following parameters: ‘-stand_call_conf 30 - stand_emit_conf 0’. Variants with no genotyped allele in any sample are removed with the GATK command SelectVariants and the parameter ‘--removeUnusedAlternates’, and variant lines where the only remaining allele is a symbolic deletion (*:DEL) are also removed using grep.

SVs were called using the svtools best practices pipeline (https://github.com/hall-lab/svtools/blob/master/Tutorial.md). First, per-sample SV calls were generated with extract-sv-reads, lumpyexpress, and svtyper using the same versions and parameters as the 14 sample analysis. Next, the calls were merged into 100-sample callsets for each pipeline using the following sequence of commands and parameters from the docker container

~~~
halllab/svtools@sha256:f2f3f9c788beb613bc26c858f897694cd6eaab450880c370bf0ef81d85bf8 d45
‘svtools lsort’
‘svtools lmerge -f 20’
‘create_coordinates’
~~~

The merged calls were then re-genotyped for each sample using the previous svtyper command. Copy number histograms were generated for each sample using the command cnvnator_wrapper.py with window size 100 (-w 100) in the docker container halllab/cnvnator@sha256:c41e9ce51183fc388ef39484cbb218f7ec2351876e5eda18b709d82b7e8af3a2. Each SV call was annotated with its copy number from the histogram file using the command ‘svtools copynumber’ in that same docker container with the parameters ‘-w 100 -c coordinates’. Finally, the per-sample genotyped and annotated VCFs were merged back together and refined with the following sequence of commands in the svtools docker container:

~~~
svtools vcfpaste
svtools afreq
svtools vcftobedpe
svtools bedpesort
svtools prune -s -d 100 -e “AF”
svtools bedpetovcf
svtools classify -a
repeatMasker.recent.lt200millidiv.LINE_SINE_SVA.GRCh38.sorted.bed.gz -m large_sample
~~~

### Cross-center variant comparisons for the 100-sample analysis

The VCF of SNPs and indels was split into per-sample VCFs using the command ‘bcftools view’ with the following parameters: ‘-a -c 1:nref’. Additionally, any remaining variant lines with only the symbolic allele (*) remaining were removed. Pairwise comparisons between the same sample processed by different pipelines were performed using hap.py using the same commands as the 14 sample analysis. Variant concordance rates per sample were calculated using results from the extended.csv output file produced by hap.py the following formula: TRUTH.TP/(TRUTH.TP + TRUTH.FN + QUERY.FP). The reported statistics were calculated using all variants genome-wide except those that were marked LowQual by GATK. No VQSR-based filtering was used. Fig 3a reports the mean rates across all 100 samples for each pairwise comparison of pipelines.

The per-pipeline SV VCFs were converted to BEDPE using the command ‘svtools vcftobedpe’ in the docker container halllab/svtools@sha256:f2f3f9c788beb613bc26c858f897694cd6eaab450880c370bf0ef81d85bf8d45. The variants were compared using bedtools pairtopair as in the 14 sample analysis. Next they were classified into “hard”, “medium”, and “easy” genomic regions by intersecting each breakpoint with BED files describing the regions using ‘bedtools pairtobed’. Variants were classified by the most difficult region that either of their breakpoints overlapped (see compare_round3_by_region.sh in https://github.com/CCDG/Pipeline-Standardization). Then, the variants were extracted and annotated in per-sample BEDPE files with the script compare_based_on_strand_output_bedpe.py (in https://github.com/CCDG/Pipeline-Standardization). The BEDPE files were converted to VCF using ‘svtools bedpetovcf’ and sorted using ‘svtools vcfsort’. The number of shared and pipeline-unique variants were counted using ‘bcftools query’ (version 1.6) to extract the genomic region and concordance status of each variant, then summarized with ‘bedtools groupby’ (v2.23.0). The rates of shared variants per sample were calculated using the output of this file with the following formula: match/(match + 0-only + 1-only).

### Mendelian error (ME) rate calculation

SNPs and indels that were classified by hap.py into categories (shared between pipelines, or unique to one pipeline) were further characterized by looking at the ME rate for each of the offspring in the trios/quads. For each offspring in the sample set, the parents and offspring sample VCFs output by hap.py were merged together using ‘bcftools merge --force-samples’ (v1.3), and the genotypes from the first pipeline in the pair were extracted. Any variants with missing genotypes or uniformly homozygous genotypes were excluded using ‘bcftools view -g ^miss’ and ‘bcftools view -g het’. A custom python script (classify_mie.py in https://github.com/CCDG/Pipeline-Standardization) was used to classify each variant as uninformative, informative with no Mendelian error, or informative with Mendelian error. Total informative error and non-error sites in each genomic region were counted for shared sites and unique sites separately, and ME rate was calculated by dividing the number of ME sites by the total number of informative sites. A similar calculation was performed for the per-sample SV VCFs produced by the SV concordance calculations. Figs 3b and S3 report the mean ME rate across 44 offspring-parent trios for each pairwise pipeline comparison.

### Variant quality evaluation

To evaluate possible causes of remaining differences between pipelines, we extracted variant quality scores for each variant type and summarized them by concordance status in each pairwise pipeline comparison across 100 samples. For SNPs and indels, the QUAL field was extracted along with the concordance annotation from the per-sample hap.py comparison VCFs using ‘bcftools query’ (version 1.6). The median QUAL score for each category was reported using ‘bedtools groupby’. For SVs, MSQ (mean sample quality) is a more informative measure of variant quality, so this field was extracted and summarized in a similar way.

---

**Supplementary Figure 1.**
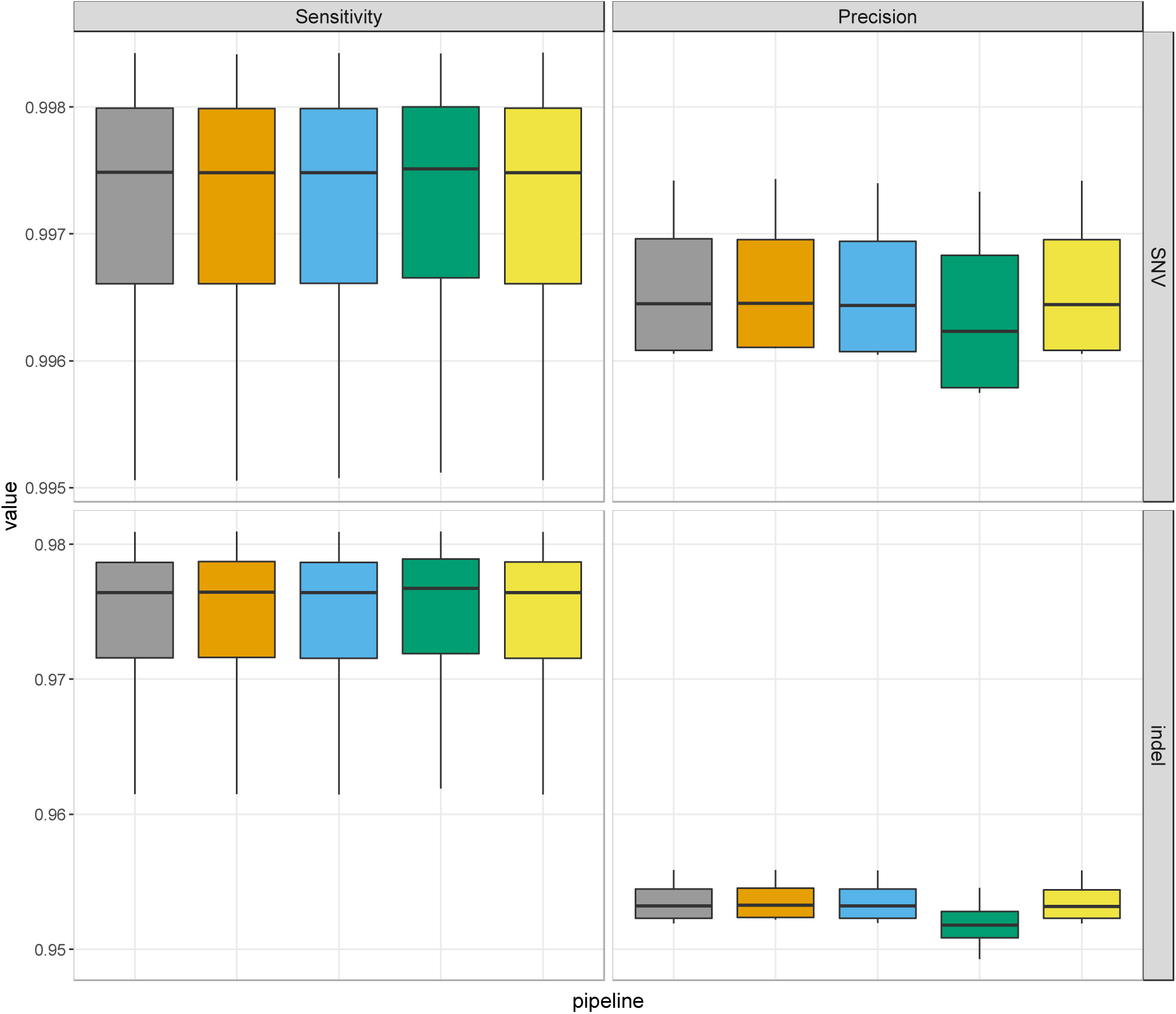
Sensitivity and precision to the GiaB gold standard variants were very similar across pipelines for all four NA12878 replicates.

**Supplementary Figure 2.**
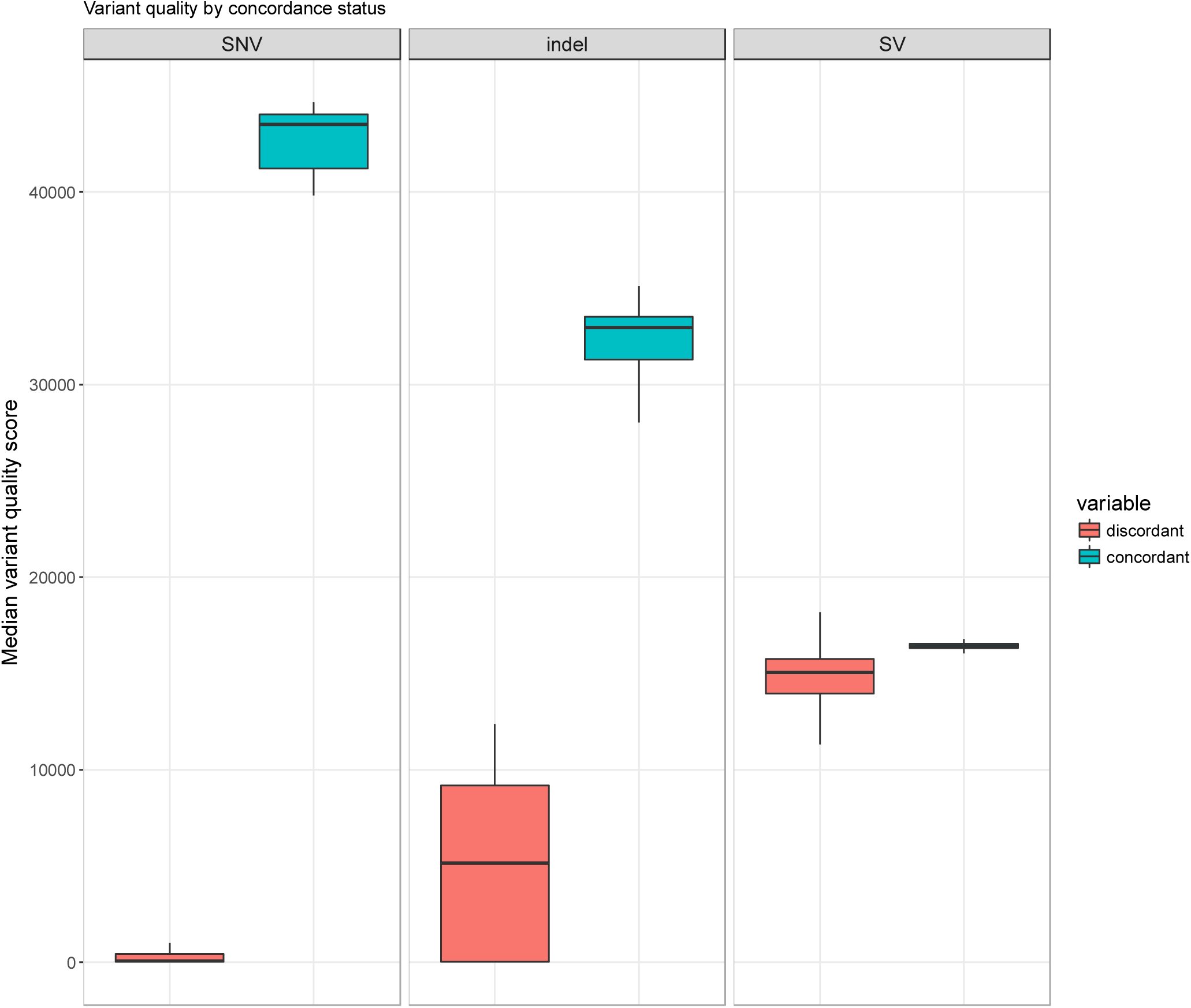
The median variant quality score (QUAL field from the GATK VCF; MSQ INFO field from the LUMPY SV VCF) was calculated for each sample, with variants partitioned by their status in each pairwise pipeline comparison.

**Supplementary Figure 3.**
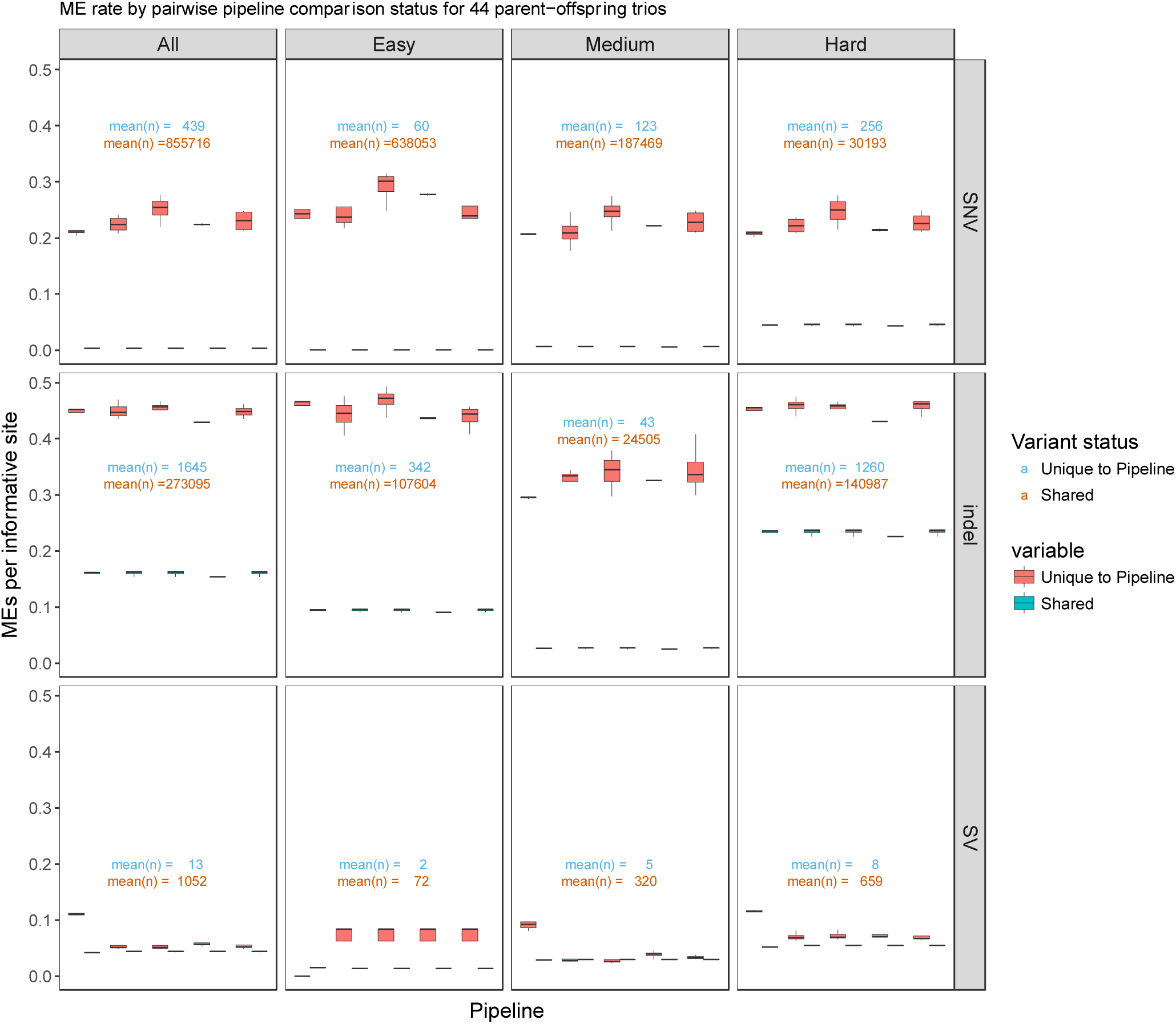
The rate of Mendelian error for each of 44 parent-offspring trios was calculated for variants shared between two pipelines as well as variants unique to one pipeline. The error rate was determined using informative sites only. In most variant types and genomic regions, variants unique to each pipeline show similar error rates, indicating that no pipeline is introducing variant calling errors or improvements in a biased way. The exception is SVs, where unique variants from one pipeline have a higher error rate than other pipelines; but, note that this is caused by a tiny number of discordant calls.

**Supplementary Table 1.**
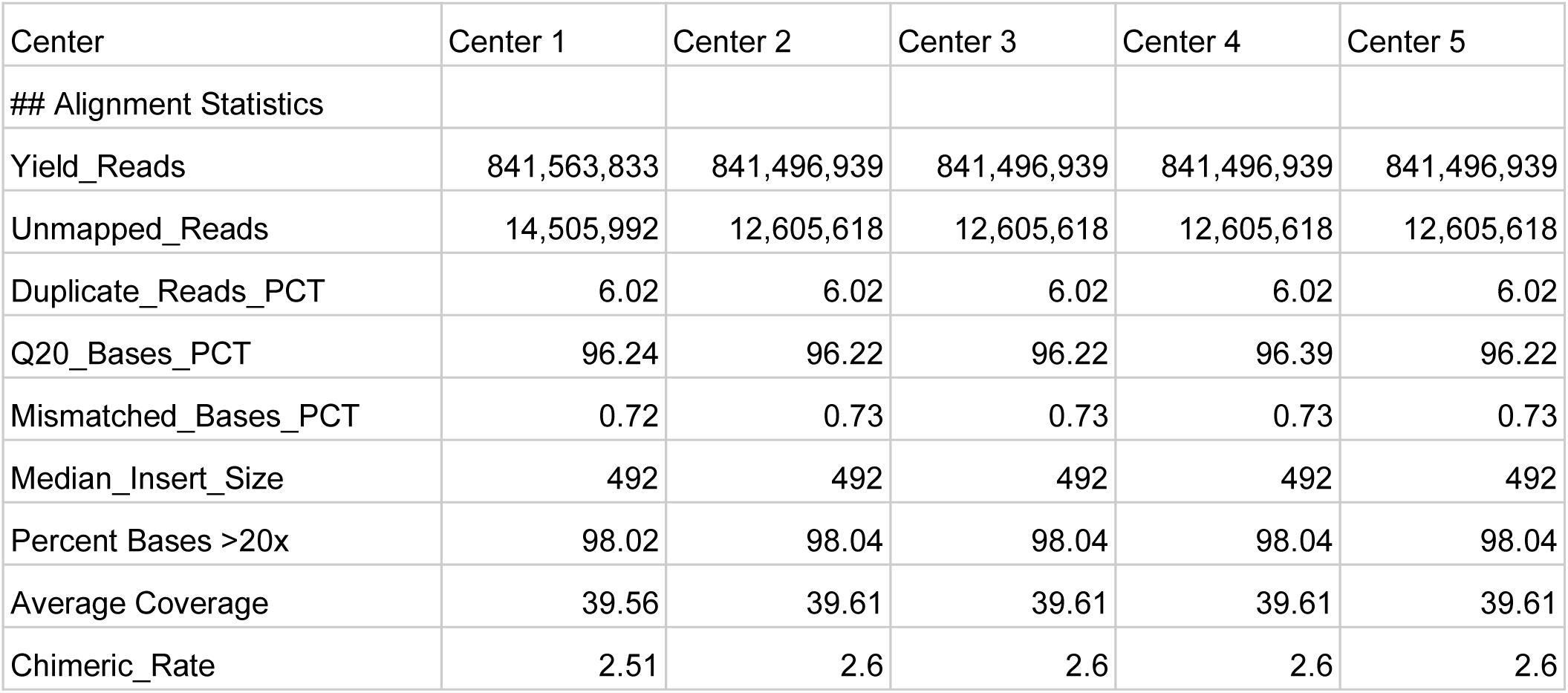
Select alignment statistics for NA19431, post-harmonization.

